# Pyoverdine-antibiotic combination treatment: its efficacy and effects on resistance evolution in *Escherichia coli*

**DOI:** 10.1101/2024.04.17.589862

**Authors:** Vera Vollenweider, Flavie Roncoroni, Rolf Kümmerli

**Affiliations:** Department of Quantitative Biomedicine, University of Zurich, Zurich, Switzerland

## Abstract

Antibiotic resistance is a growing concern for global health, demanding innovative and effective strategies to combat pathogenic bacteria. Pyoverdines, iron-chelating siderophores produced by environmental *Pseudomonas* spp., present a novel promising approach to induce growth arrest in pathogens through iron starvation. While we have previously demonstrated the efficacy of pyoverdines as antibacterials, our understanding of how these molecules interact with antibiotics and impact resistance evolution remains unknown. Here, we investigate the propensity of different *Escherichia coli* variants to evolve resistance against pyoverdine, the cephalosporin antibiotic ceftazidime, and their combination. We found that strong resistance against ceftazidime and weak resistance against pyoverdine evolved in the wildtype *E. coli* strain under single and combination treatment. Ceftazidime resistance was linked to mutations in outer membrane porin genes (*envZ* and *ompF*), whereas pyoverdine resistance was associated with mutations in the oligopeptide permease (*opp*) operon. In contrast, resistance phenotypes were attenuated under combination treatment, particularly in an *E. coli* strain carrying a costly multicopy plasmid. Altogether, our results show that pyoverdine as an antibacterial is particularly potent and evolutionarily robust against plasmid-carrying *E. coli* strains, presumably because iron starvation compromises both cellular metabolism and plasmid replication.

## Introduction

Antibiotic-resistant bacteria pose a significant global health problem with estimates predicting 10 million annual deaths attributable to AMR (antimicrobial resistance) by 2050^1,2^. Consequently, there is an urgent need for new and more sustainable approaches to effectively control pathogens and to counteract the emergence and spread of resistance^3–5^. While several alternative approaches to antibiotics are under investigation (e.g., phage therapy, antibacterial peptides and anti-virulence approaches)^6–11^, we have previously shown that siderophores from environmental bacteria can have strong inhibitory effects on opportunistic human pathogens through the induction of iron limitation^12^.

Particularly, we found that pyoverdines from environmental *Pseudomonas* spp. show a great structural diversity and high iron-chelation properties and are thus able to induce iron starvation and growth arrest in difficult-to-treat pathogens such as *Acinetobacter baumannii* and *Staphylococcus aureus*^12^. Furthermore, we observed that pyoverdine treatment improved the survival of infected *Galleria mellonella* host larvae, while having minimal negative effects on mammalian cell lines and erythrocytes. Finally, we found low levels of resistance emerging in pathogens exposed to the pyoverdine treatment compared to a conventional antibiotic.

While pyoverdines (or synthesized derivates) could become potential novel antibacterials, one important aspect that needs closer examination is how pyoverdine treatment interacts with antibiotics and how combination treatment affects selection for antibiotic resistance. Combination treatments are well-established in clinical settings and have been proven successful against *Mycobacterium tuberculosis* infections^13,14^, HIV infections^15,16^ and *Plasmodium falciparum* malaria infections^17^. Here, we explored the efficacy and evolutionarily sustainability of combination therapy involving pyoverdine by using experimental evolution with the model pathogen *Escherichia coli*. Specifically, we subjected this pathogen to either pyoverdine 3G07 (one of the most potent pyoverdines against a range of pathogens^12^; Fig. S1), ceftazidime (a cephalosporin antibiotic) or a combination of the two.

Additionally, we asked whether the genetic background of the pathogen influences the selection for resistance. We were particularly interested in whether the carriage of elements conferring resistance to other antibiotics affects resistance evolution against pyoverdine and/or ceftazidime. To address this question, we used three different *E. coli* strains^18^. We took wild-type *E. coli* MG1655 (MG) as the naive susceptible strain. We further used two isogenic MG variants that both carried the β-lactamase gene *bla*_TEM-1_, conferring resistance to β-lactams, such as ampicillin, but not against ceftazidime. One of the strains harboured the *bla*_TEM-1_ resistance gene on the chromosome (MGc), while the other one carried the same gene on a small multicopy plasmid (MGp). Carrying resistance elements is typically associated with fitness costs and the relative costs of resistance may differ between chromosomal and plasmid-based variants. Plasmids can be costly in the absence of antibiotics because they often contain many genes beyond those involved in resistance, including genes for plasmid replication and maintenance^19,20^. Chromosomally integrated resistance elements should be less costly as long as they do not affect the expression of other genes encoding critical cellular functions^21,22^. Here, we test whether these differential costs affect the evolution of resistance against pyoverdine 3G07 and ceftazidime single and combo treatments.

Before conducting the experimental evolution experiment, we established dose-response curves for the three *E. coli* strains when exposed to pyoverdine 3G07, ceftazidime, and the combo treatment. This allowed us to define drug efficiency ranges and drug interactions. During the experimental evolution experiment, we cultured the three *E. coli* strains in four conditions: pyoverdine 3G07 and ceftazidime single treatments, the combo treatment, and media without treatment. Each condition was replicated six times and cultures were daily transferred to fresh media. Following experimental evolution, we screened evolved populations for resistance phenotypes and sequenced a subset of populations to uncover the genetic basis of putative resistance mechanisms.

## Results

### Pyoverdine and ceftazidime treatments curb the growth of all *E. coli* strains

To confirm the efficacy of pyoverdine 3G07 and ceftazidime against *E. coli* MG1655, we exposed the three MG strains to increasing drug concentrations and assessed their growth performance (Fig. 1). For both treatments we observed conventional dose-response curves, with no growth inhibition at low concentrations, followed by a decrease in bacterial growth at intermediate and high concentrations. The dose-response curves were similar for all three MG strains, with one exception: MGc showed a slightly shallower response to pyoverdine than the other two strains. For pyoverdine, we estimated the following half maximal inhibitory (IC50) concentrations: MG = 0.15, MGc = 0.27, MGp = 0.17. Note that the absolute pyoverdine concentration cannot be assessed from the crude extracts we used here, and that is why we express concentrations relative to the weighed amount of 6 mg/mL. For ceftazidime, estimates of IC50 concentrations were 0.31 mg/L for MG, 0.32 mg/L for MGc, and 0.29 for MGp, and were highly consistent across the three strains.

**Figure 1.**
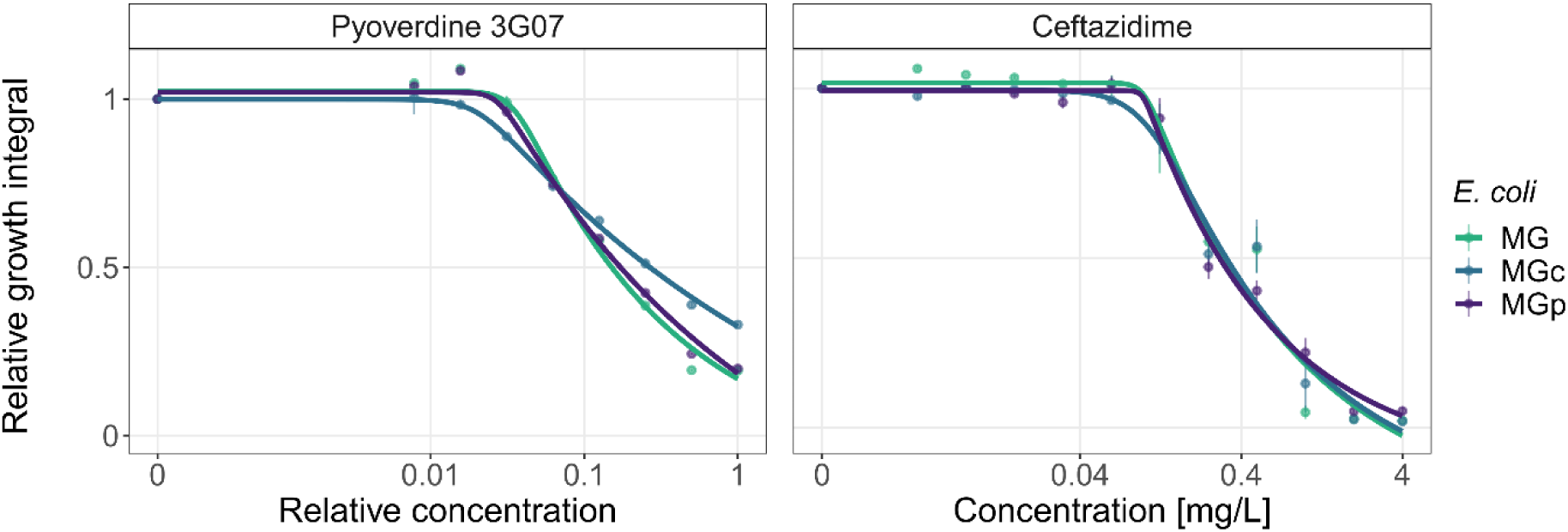
Dose-response curves for *E. coli* strains MG, MGc and MGp following treatment with pyoverdine 3G07 or ceftazidime. We exposed the three strains to increasing concentrations of pyoverdine 3G07 and ceftazidime and calculated the integral below the growth curves. These values were then scaled relative to the untreated control in CAA medium. Dots and error bars show mean values and standard errors, respectively, across a minimum of six replicates per concentration. Dose-response curves were fitted using 5-parameter logistic regressions.

### Pyoverdine 3G07 and ceftazidime show a neutral interaction

To quantify whether the interaction between pyoverdine 3G07 and ceftazidime is antagonistic, neutral (additive) or synergistic, we used the Bliss model to compare whether the combo treatment is less, equally or more inhibitory than expected, respectively^23^. For this comparison, we used a relative pyoverdine concentration of 0.16 and a ceftazidime concentration of 0.4 mg/L for both single and combo treatments, which displayed intermediate inhibition (Fig. 1). We found that all interaction terms are approximately zero, and when combined across strains, the interaction was not significantly different from the expected value (Tab. 1, t_8_ = 0.52, p = 0.6166). This result suggests that the interaction between pyoverdine 3G07 and ceftazidime is neutral (additive).

**Table 1.** Drug interaction in pyoverdine-ceftazidime combination treatment.

| Strain | Mean degree of synergy | St. error | t value | df | p value |
| --- | --- | --- | --- | --- | --- |
| MG | 0.0294 | 0.0739 | 0.3979 | 2 | 0.7291 |
| MGc | -0.0277 | 0.0574 | -0.4823 | 2 | 0.6772 |
| MGp | 0.0446 | 0.0186 | 2.4011 | 2 | 0.1383 |
| Combined | 0.0155 | 0.0297 | 0.5209 | 8 | 0.6166 |

### Plasmid-based antibiotic resistance entails a fitness cost

Next, we investigated whether carrying a plasmid encoding a resistance gene is associated with fitness costs. To this end, we competed the plasmid-carrying strain MGp against the isogenic strain MG and the chromosomally resistant strain MGc both in medium without and with ampicillin, the antibiotic to which *bla*_TEM-1_ confers resistance. For all competitions, we used 0.5 mg/L ampicillin, which represents the IC50 concentration of strain MG (Fig. S2).

In the absence of ampicillin, the fitness of MGp was significantly reduced compared to MG (one-sample *t*-test, *t*_5_ = −3.15, p = 0.0254) and MGc (*t*_5_ = −3.96, p = 0.0107) (Fig. 2). These results demonstrate that plasmid carriage is associated with fitness costs in the absence of antibiotics. Meanwhile, in the presence of ampicillin, the benefits of carrying a resistance plasmid exceeded its cost, resulting in MGp exhibiting significantly higher fitness compared to the susceptible strain MG (*t*_5_ = 29.50, p < 0.0001). Nonetheless, even in the presence of ampicillin, MGp exhibited reduced fitness compared to MGc (*t*_5_ = −2.72, p = 0.0416), showing that plasmid-based resistance entails higher costs than chromosomal-based resistance in our system.

**Figure 2.**
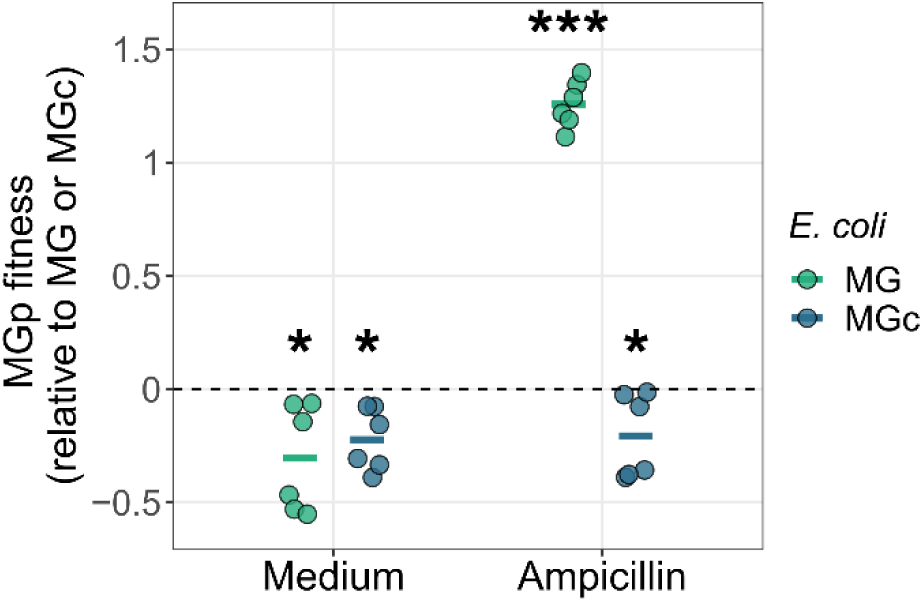
Relative fitness of plasmid-carrying *E. coli* MGp in competition against *E. coli* MG and MGc. We tested whether carrying an antibiotic resistance plasmid has a cost compared to a naive strain (MG) and a strain carrying the resistance gene on the chromosome (MGc). *E. coli* MGp was therefore competed against MG and MGc for 48 hours, starting at a 1:1 ratio. The dashed line indicates fitness parity, at which neither of the competing strains has a fitness advantage. Without any antibiotic treatment (medium), MGp had a fitness disadvantage (fitness values < 0) compared to MG and MGc, showing the cost of plasmid-based resistance. When treated with ampicillin, MGp experienced highly significant fitness advantages (fitness values > 0) compared to MG, and slight significant fitness disadvantages compared to MGc. All data are shown as means ± standard errors across six replicates from two independent experiments. Significance levels are based on one-sample *t*-tests (comparison against the null-line): * p < 0.05; ** p < 0.01; *** p < 0.001.

### Combination treatment reduces selection for resistant phenotypes in *E. coli* MGp

We then used experimental evolution to assess whether *E. coli* strains MG, MGc and MGp evolve resistance to either pyoverdine 3G07, ceftazidime or the combo treatment. We propagated six independent populations per condition and strain every other day to fresh medium for 15 transfers (30 days in total). To control for medium adaptation, we also evolved the three *E. coli* strains in the absence of drugs, resulting in a total of 72 evolving populations (Fig. S3). Subsequently, we subjected the evolved populations from the final transfer to the conditions they evolved in and compared their growth relative to the ancestral wildtypes (Fig. S4) by calculating the difference in growth between the evolved and ancestral populations.

We found that evolved MG populations grew significantly better under pyoverdine, ceftazidime and the combo treatment compared to the control populations that evolved without antibacterials (global analysis: ANOVA, F_3,20_ = 51.07, p < 0.0001, Fig. 3). This suggests that MG populations evolved (at least partial) resistance to pyoverdine, ceftazidime and the combo treatment. In contrast, evolved MGc populations only showed significantly improved growth under ceftazidime and the combo treatment but not under pyoverdine single treatment (compared to evolved control: ANOVA, F_3,20_ = 19.6, p < 0.0001). Finally, the evolved MGp populations grew significantly better only under ceftazidime treatment, but not under pyoverdine or the combo treatment (compared to evolved control: ANOVA, F_3,20_ = 30.14, p < 0.0001). These results suggest that MGp has evolved resistance to ceftazidime, but not against treatments containing pyoverdine. Taken together, our analyses suggest that the genetic background of the strains influences resistance evolution, and that resistance evolution seems lowest in MGp.

**Figure 3.**
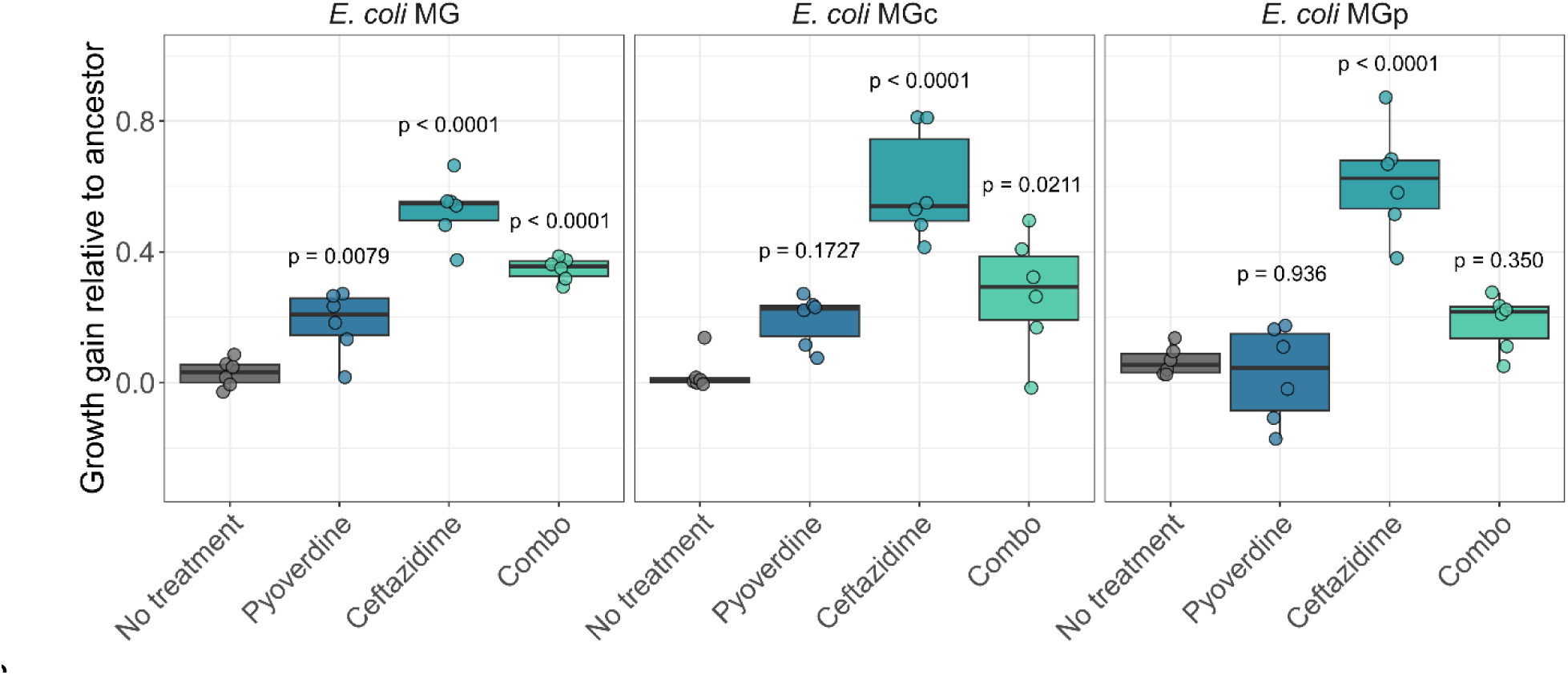
Growth gain of evolved *E. coli* MG, MGc and MGp populations relative to ancestral populations in the no-treatment control, or the three antibacterial treatments pyoverdine 3G07, ceftazidime and their combo. We exposed the evolved *E. coli* populations to the conditions in which they evolved in (no-treatment control, pyoverdine 3G07, ceftazidime and combo treatment). We further exposed the ancestors to the same four conditions. We quantified growth by calculating the integrals under the growth curves, which were then scaled relative to the ancestral growth in plain CAA medium. The difference in growth values was subsequently calculated between evolved and ancestor populations that experienced the same treatment condition. This difference corresponds to the growth gain of evolved populations. Each dot denotes a population and represents the mean value across six replicates. Box plots show the median together with the first and third quartiles and whiskers represent the 1.5x interquartile range. The p-values above antibacterial treatment boxplots indicate the significance levels relative to the no-treatment control (assessed by ANOVA with adjusted p-values using the Tukey HSD method, α = 0.05).

### Pyoverdine 3G07 masks rather than prevents ceftazidime resistant phenotypes in *E. coli* MGp populations

Given the reduced growth gain in evolved *E. coli* MGp populations in the combo treatment compared to the ceftazidime single treatment, we wondered whether pyoverdine 3G07 can indeed reduce selection for resistance or whether it simply masks the full ceftazidime resistant phenotype. To differentiate between these two possibilities, we exposed the MGp ancestor and MGp populations evolved in the combo treatment to single pyoverdine 3G07 and ceftazidime treatments as well as the no-drug control treatment. We monitored their growth for 48 hours (Fig. S5) and calculated the growth gain as described above.

We found that the MGp populations evolved in the combo treatment showed significant differences in growth gain when exposed to single treatments (ANOVA, F_2,15_ = 32.90, p < 0.0001, Fig. 4). No significant difference in growth gain occurred between the no-treatment control and the pyoverdine treatment (Tukey post-hoc: p = 0.6635), suggesting low levels of pyoverdine resistance. By contrast, growth gains in MGp populations evolved in the combo treatment were substantial under ceftazidime exposure and significantly higher in the ceftazidime single (mean±SD: 0.392±0.082) than in the combo (0.283±0.056) treatment (p = 0.0207). This suggests that pyoverdine 3G07 does not prevent the evolution of ceftazidime resistant phenotypes in MGp populations under combo treatment, but rather masks them.

**Figure 4.**
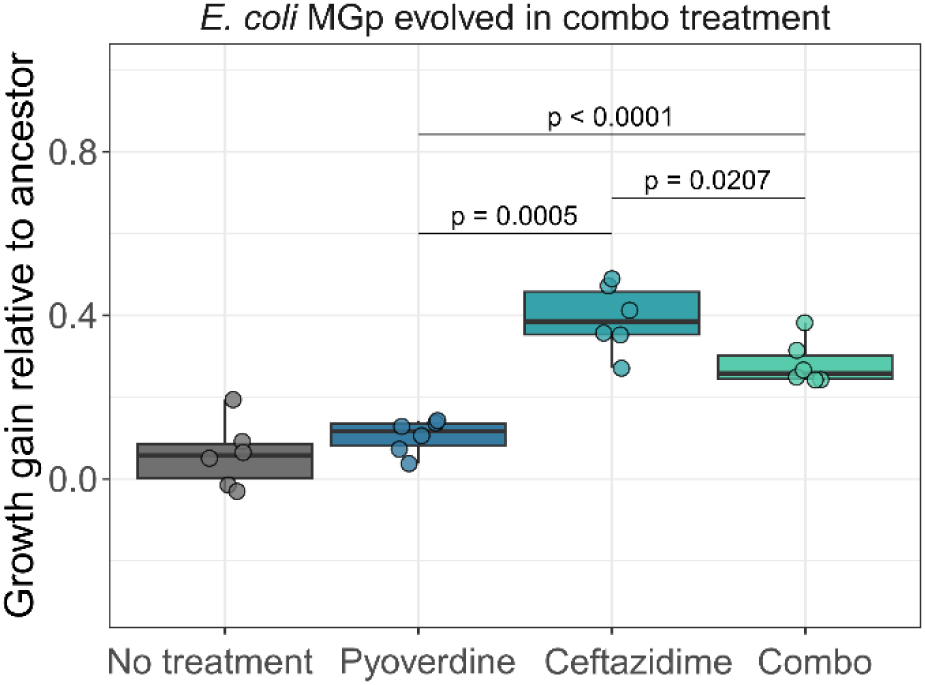
Growth gain of *E. coli* MGp populations evolved in the combo treatment and tested under single pyoverdine 3G07 and ceftazidime treatments. We exposed the six *E. coli* MGp populations that evolved in the combo treatment to single pyoverdine and ceftazidime treatments, as well as to the no-treatment control and the combo treatment in which they evolved. Growth gain is represented by the scaled difference in growth integral between evolved and ancestor populations. Each dot denotes a population and represents the mean value across six replicates. Box plots show the median along with the first, while whiskers represent the 1.5x interquartile range. Statistical analysis was performed using ANOVA with adjusted p-values using the Tukey HSD method (α = 0.05).

### Mutation frequencies of evolved *E. coli* MG and MGp populations

The above growth screen suggests that *E. coli* MG populations developed resistance to both pyoverdine 3G07 and ceftazidime treatments, while *E. coli* MGp specifically acquired resistance to ceftazidime. To link these observed phenotypes to mutational patterns, we sequenced four evolved populations per antibacterial treatment and three no-treatment control populations from the final day of experimental evolution. Overall, we sequenced 30 populations, 15 each for MG and MGp. We excluded mutations present in ancestral clones and focussed on those accumulated during experimental evolution.

In total, we identified 453 and 1221 mutations across all MG and MGp populations, respectively. Mutation frequencies within populations ranged from 5% (our lower cut-off) to 100%. Across all evolution conditions and strains, mutation frequencies exhibited a bimodal distribution: the majority (1371) of mutations occurred at low frequencies (< 30%), while a minority (303) occurred at higher frequencies (> 30%; Fig. 5A). The low-frequency mutations appeared in intergenic regions (47.8%), pseudogenes (24.9%), coding regions (22.1%), and noncoding regions (5.1%; Tab. S1). We excluded these low-frequency mutations from further analyses since we argue that many of them might be neutral and subject to drift rather than selection.

**Figure 5.**
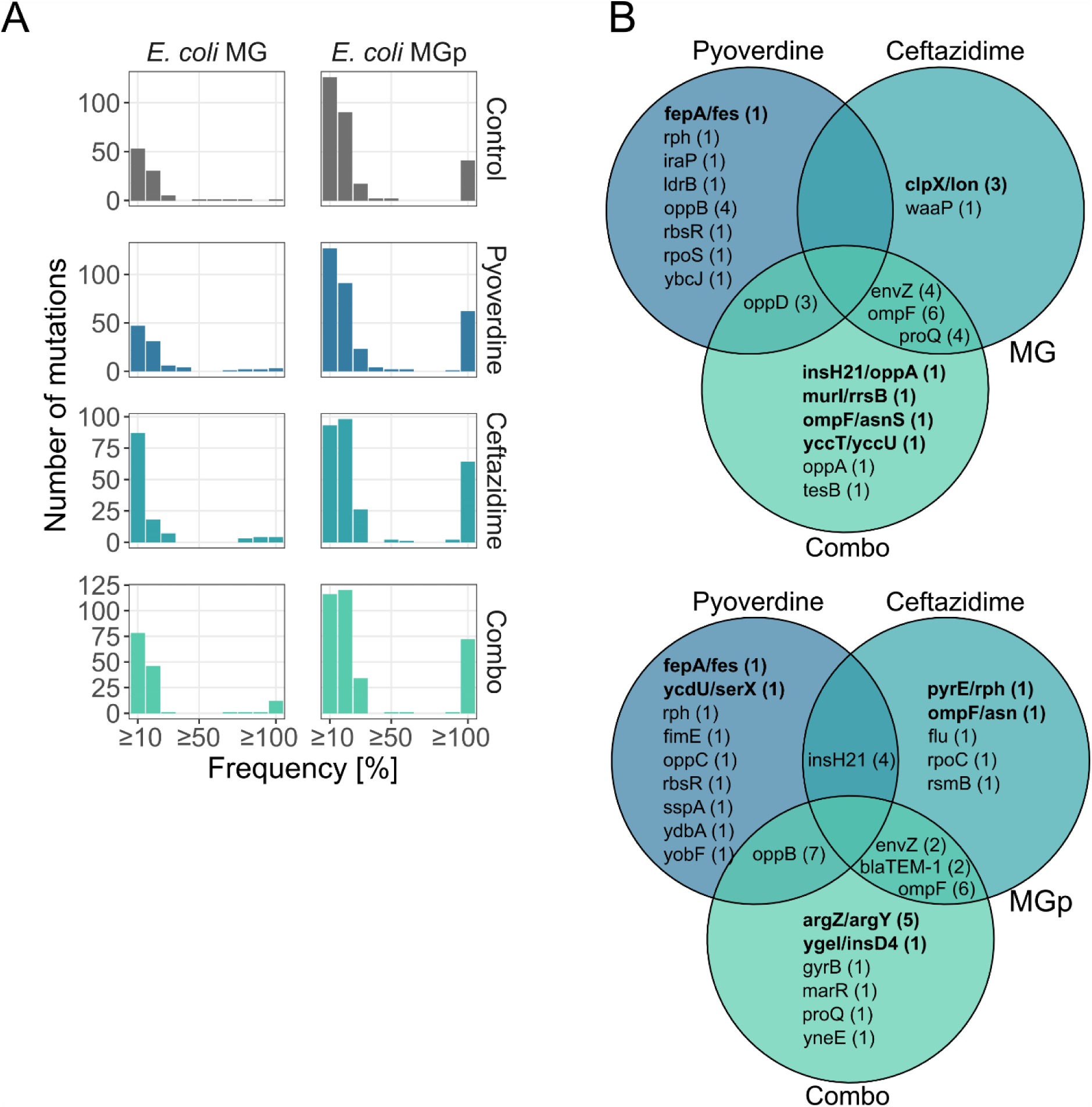
Mutational patterns in *E. coli* MG and MGp after evolution in pyoverdine 3G07, ceftazidime or the combo treatment. (A) Bimodal distribution of the number of mutations per population of MG and MGp after evolution. In total, 1371 mutations were present at low frequency (< 30%), while 303 mutations were present at higher frequency (> 30%). (B) Venn diagram showing the genes unique to or shared between populations evolved in pyoverdine 3G07, ceftazidime or the combo treatment for MG and MGp. Bold letters indicate mutations in intergenic regions and numbers in brackets indicate the number of overall mutations within that region/gene.

Focusing on high-frequency mutations (> 30%), we identified single-nucleotide polymorphisms (SNPs) as the most common mutation (135), followed by deletions (97), insertions (50), mobile element insertions (MOB; 20), and amplification (1). Most mutations appeared in intergenic regions (52.5%), followed by coding regions (47.2%) and pseudogenes (0.3%).

Next, we excluded genes with synonymous mutations and genes that mutated in both no-treatment control and antibacterial treatments as they are probably associated with medium adaptation. After this filtering process, the frequencies of the remaining mutations were similar in the pyoverdine treatment (12 and 15 mutations), the ceftazidime treatment (11 and 12), and the combo treatment (15 and 19 mutations in MG and MGp, respectively), and did not differ between MG and MGp (Fisher’s exact test, p = 0.9619).

### Comparison of mutational patterns in single and combo treatments

To identify putative key mutations responsible for resistance phenotypes, we compared the number of mutations unique to a particular antibacterial treatment with those shared between treatments. This analysis yielded little overlap in the mutational patterns between treatments (pairwise comparisons) and no mutation occurred in all three treatments (Fig. 5B). However, those mutations that occurred in two treatments were frequent and surfaced in multiple populations. We now go through the list of these mutations and assess their potential association with antibacterial resistance.

For ceftazidime single and combo treatments, mutations in the following genes occurred at least twice: *ompF* (MG = 6/ MGp = 6), *envZ* (4/2), *bla_TEM-1_* (0/2), *proQ* (4/0). *OmpF* encodes the outer membrane porin F^24,25^, a known entry point for antibiotics. Loss-of-function mutations in *ompF* reduce membrane permeability, leading to increased levels of antibiotic resistance^26,27^. Mutations in *envZ* could have a similar effect. EnvZ is a periplasmic protein that senses changes in the environment and controls the phosphorylation state of its cognate response regulator OmpR, which is a repressor of *ompF*. Thus, mutations in *envZ* could result in the down-regulation of *ompF* expression^24,25^. *Bla_TEM-1_* is the resistance gene introduced in *E. coli* MGp^18^ and encodes a beta-lactamase with high activity against ampicillin. In its original form, it has minimal activity against ceftazidime, but mutations can expand its activity range to ceftazidime^28–30^. ProQ is an RNA chaperon involved in the posttranscriptional control of ProP levels, which are plasma membrane transporters that sense and respond to osmotic changes ^31^. Here, it is less clear how mutations in this gene could be involved in ceftazidime resistance.

For the pyoverdine 3G07 single treatment, we identified a SNP in the regulatory region between genes encoding the ferric enterobactin outer membrane transporter FepA^32–34^ and the ferric enterobactin esterase Fes^35,36^ in both MG and MGp (Fig. S6). Enterobactin, a high-affinity siderophore produced by *E. coli*, binds iron and is transported by FepA, while Fes catalyses the hydrolysis of the ferric-enterobactin complex in the cytosol^37^. Notably, the ferric uptake regulator (Fur) protein binding site, which is crucial for regulating the response to iron starvation, is situated between FepA and Fes^38–40^. Consequently, mutations in this region may result in a constant dissociation of Fur and consequently to a constitutive expression of genes related to enterobactin transport and utilization. The upregulation of enterobactin could be beneficial, as this siderophore has higher iron affinity than pyoverdine and could thus alleviate the iron starvation imposed by pyoverdine.

Additionally, we found that mutations in the genes of the *opp*-operon were common (MG/MGp: *oppA* = 2/0, *oppB* = 4/7, *oppC* = 0/1, *oppD* = 3/0) in populations of the pyoverdine single and combo treatments, reaching high frequencies (mean±SE, MG: 0.83±0.05; MGp: 0.80±0.08). This operon encodes an oligopeptide importer^41^ and the uncovered mutations are likely associated with a loss-of-function (Fig. S7, Tab. S2). Since pyoverdine is a modified oligopeptide, we speculate that loss-of-function mutations in the *opp*-operon could potentially prevent the uptake of apo-pyoverdine. A reduction in apo-pyoverdine uptake could potentially prevent this molecule from interfering with intracellular iron homeostasis.

### Enhanced growth of *E. coli ΔoppA* and *ΔoppB* mutants in pyoverdine 3G07 treatment

While our previous work yielded little evidence for resistance evolution against pyoverdine in other pathogens (*Acinetobacter baumannii*, *Klebsiella pneumoniae*, *Staphylococcus aureus*)^12^, we here identify the *opp* operon as a potential mutational target conferring a moderate level of pyoverdine resistance. The operon *oppABCDF* consists of five genes, encoding a high affinity oligopeptide ABC transporter system with broad specificity^42,43^. *OppA* encodes a periplasmic binding protein, which interacts with the two inner membrane subunits o*ppBC*^44,45^, and the ATP binding subunits *oppDF*^41,46^.

To understand whether mutations in the *oppABCDF* operon indeed confer resistance to pyoverdine, we subjected the *E. coli* mutants Δ*oppA*, Δ*oppB*, Δ*oppC* and Δ*oppD* to pyoverdine 3G07 treatment and compared their growth to that of the parental wildtype BW25133. These mutants originate from the Keio collection^47^, where the gene of interest is replaced with a kanamycin resistance cassette and thus allows to analyse the effects of complete loss of gene function.

We first examined whether the lack of a functional Opp-transporter has fitness consequences for *E. coli* in the absence of antibacterial treatment (Fig. 6; grey growth kinetics and circles). We found that growth was reduced in Δ*oppA* (two-sample *t*-test: *t*_10_ = 2.46, p = 0.0338) and Δ*oppB* (*t*_10_ = 6.34, p = 0.0001) compared to wildtype but not in Δ*oppC* (*t*_10_ = 0.36, p = 0.7253) and Δ*oppD* (*t*_10_ = 0.98, p = 0.3512). Overall, growth effects were small suggesting that the Opp-transporter is not essential for *E. coli*.

**Figure 6.**
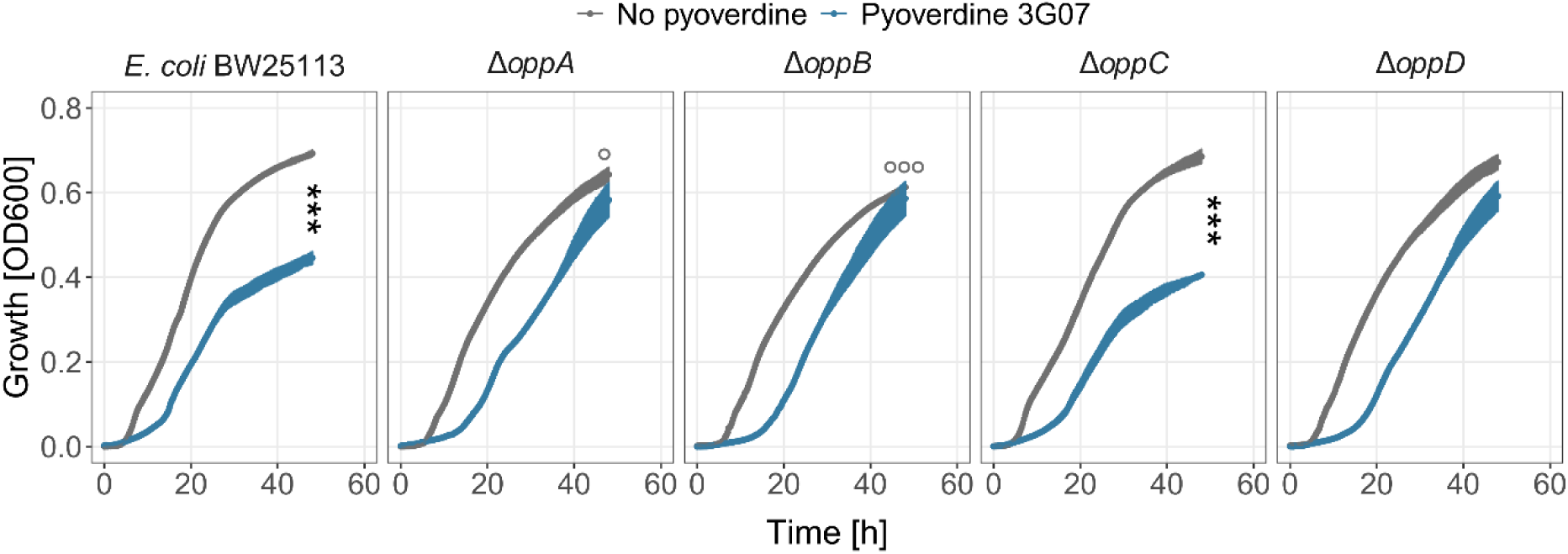
Growth of *E. coli opp* single-gene knockout mutants in the presence and absence of pyoverdine 3G07. Growth kinetics of the *E. coli* knockout mutants Δ*oppA*, Δ*oppB*, Δ*oppC*, Δ*oppD* and the *E. coli* wildtype BW25113 in medium without pyoverdine (grey lines) and with pyoverdine (blue lines) treatment over 48 hours. Dots and error bars show mean values and standard errors, respectively, across a minimum of six replicates per treatment from two independent experiments. Significance levels are based on two-sample *t*-tests with α < 0.05. Circles represent difference in the end-point growth between the *E. coli* wildtype and the mutants in the no-pyoverdine control (grey lines). Asterisks denote significant differences in the end-point growth between the no-pyoverdine condition and the pyoverdine treatment in each panel.

Next, we tested whether the lack of specific Opp-proteins alleviates the significant growth reduction that pyoverdine treatment has on the wildtype (*t*_10_ = −13.76, p < 0.0001, Fig. 6; blue growth kinetics and asterisk). Indeed, we found that growth yield after 48 hours was restored to the level of the no-pyoverdine treatment control in Δ*oppA* (*t*_10_ = - 1.33, p = 0.2146) and Δ*oppB* (*t*_10_ = −0.61, p = 0.5676), and to some extent in Δ*oppD* (*t*_10_ = - 1.97, p = 0.0777), but not in Δ*oppC* (*t*_10_ = −15.06, p < 0.0001). However, albeit achieving the same yield, the growth of all *opp*-mutants was still substantially slowed down compared to the untreated controls during the early growth phase. These results indicate that mutations in the *opp* operon can confer partial resistance to pyoverdine.

## Discussion

We previously demonstrated the potent inhibitory effects of iron-scavenging pyoverdines from environmental *Pseudomonas* spp. on opportunistic human pathogens through the induction of iron limitation^12^. Building on this work, we here investigated the interaction between pyoverdine 3G07 and the antibiotic ceftazidime to explore how combination therapy affects resistance evolution in three *E. coli* strains. We found neutral drug interactions between pyoverdine and ceftazidime and observed that pyoverdine could not prevent the evolution of ceftazidime resistance in a combination treatment in the naive *E. coli* strain MG during experimental evolution. In contrast, the presence of pyoverdine reduced ceftazidime resistant phenotypes in *E. coli* MGc (mild effect) and *E. coli* MGp (strong effect), two strains carrying the β-lactamase gene *bla*_TEM-1_ (conferring resistance to ampicillin) on the chromosome and a small multicopy plasmid, respectively. Genetic analyses revealed that mutations in the outer membrane porin F (OmpF) and the sensor histidine kinase EnvZ were associated with ceftazidime resistance. Curiously, the frequency of these mutations was similar in both *E. coli* strains MG and MGp, regardless of whether they were treated with ceftazidime alone or in combination with pyoverdine. This suggests that pyoverdine cannot prevent resistance evolution per se but can effectively mask resistant phenotypes.

We found additive (neutral) effects between pyoverdine 3G07 and ceftazidime. Given their independent modes of action – pyoverdine induces iron starvation, while ceftazidime causes cell lysis – neutral interactions between these two classes of antibacterials can be anticipated. Neutral drug interactions are beneficial from a clinical perspective because they reduce the risk that a single mutation can simultaneously confer resistance to both antibacterials^48^. This notion is supported by our data showing that there is hardly any overlap in high-frequency mutations between the pyoverdine and ceftazidime single treatments (Fig. 5). However, neutral drug interactions are not necessarily making treatments evolutionarily more sustainable, as the combined drugs should not influence each other’s resistance evolution^49,50^. This notion is also supported by our data, as we found (i) similar mutational patterns in populations subjected to ceftazidime single and combo treatments (Fig. 5), and (ii) that ceftazidime resistance was unmasked when pyoverdine was omitted from the combo treatment (Fig. 4). Thus, our results show that pyoverdine cannot reduce selection for ceftazidime resistance, but it can safely be combined with it as cross-resistance is unlikely to evolve.

An important question to address is why pyoverdine is particularly potent at masking the ceftazidime resistance in the plasmid-carrying *E. coli* MGp strain, but not in the naive *E. coli* MG strain. We propose that a dual metabolic burden is responsible for this effect. For one thing, MGp requires extra resources for plasmid maintenance and replication. At the same time, pyoverdine induces iron starvation with iron being an essential trace element required for DNA replication^51,52^. Hence, the dual metabolic burden could explain why evolved *E. coli* MGp populations remained more susceptible to the combo treatment than evolved *E. coli* MG populations, even though the two evolved strains exhibited similar mutational patterns regarding pyoverdine and ceftazidime resistance. While we used a costly multicopy plasmid containing the *bla*_TEM-1_ gene for our experiments, we anticipate that the dual metabolic burden applies to any costly plasmid. Overall, our findings indicate that targeting iron metabolism seems to be particularly effective for pathogens with high iron requirements.

While our previous work yielded little evidence for pyoverdine resistance evolution in several pathogens (*A. baumannii*, *S. aureus*, *K. pneumoniae*, *P. aeruginosa*)^12^, we here identified a first putative target involved in partial pyoverdine resistance in *E. coli*. We found that loss-of-function mutations in the *oppABCDF* operon reduced susceptibility to pyoverdine (Fig. 6). The *oppABCDF* operon codes for a periplasmic oligopeptide importer^42^. Discovering this target of evolution surprised us because it contradicts the basic premise that pyoverdine can solely inhibit pathogens by sequestering iron extracellularly^12^. Accordingly, we predicted that mechanisms reducing drug entry into the cell should not be an effective resistance mechanism against pyoverdine. Now we find exactly such a mechanism to be associated with pyoverdine resistance. To reconcile this apparent discrepancy, we propose a dual mode of action of how pyoverdine can inhibit *E. coli*. The first mode of action occurs extracellularly, whereby pyoverdine chelates iron in the environment, thereby withholding it from *E. coli*. The second mode of action could involve the translocation of iron-free (apo-) pyoverdine via non-specific importers (e.g., porins) to the periplasm and from there via non-specific oligopeptide transporters (e.g. Opp-transporter) to the cytosol. In the cytosol, apo-pyoverdine can potentially interfere with cellular iron homeostasis. Pyoverdine is an oligopeptide (Fig. S1) and previous studies showed that the Opp transporter is non-selective towards amino acid side chains, allowing the transport of peptides of various lengths and structures^45^. Loss-of-function mutations in any of the *opp* genes could thus potentially block this second mode of action. While novel and speculative at the same time, our two modes of action model would explain why mutations in the *opp*-operon (cutting the second mode of action) only leads to partial pyoverdine resistance, given that the first mode of action remains functional. Clearly, further experiments are required to ascertain whether apo-pyoverdine is indeed translocated into *E. coli* cells and whether this mechanism is specific to *E. coli*, given the absence of similar mutational targets in other pathogens^12^.

In summary, we show that pyoverdine is a potent antibacterial against *E. coli* both as single treatment and in combination with the antibiotic ceftazidime. The interaction between pyoverdine and ceftazidime is neutral, both in terms of treatment efficacy and selection of resistance evolution. While pyoverdine is evolutionary more robust than ceftazidime, pyoverdine as an adjuvant only masks but does not prevent the evolution of ceftazidime resistance. Pyoverdine treatment is particularly potent against plasmid-carrying *E. coli* strains, possibly due to a dual metabolic burden associated with plasmid maintenance. This finding suggests the pyoverdine treatment might be particularly effective against pathogens harbouring costly antibiotic-resistance plasmids.

## Acknowledgments

We thank Richard Allen and Clémentine Laffont for insightful discussion. This work was supported by funding from the Swiss National Science Foundation (grant number 31003A_182499) and the University Research Priority Program (URPP) «Evolution in Action».

## Materials and methods

### Bacterial strains

We used the *E. coli* strain MG1655 (MG) and two isogenic constructs containing the β- lactamase gene *bla*_TEM-1_ either on the chromosome (MGc) or on a non-transmissible multicopy plasmid (MGp) with identical promoters. TEM-1 confers resistance to ampicillin and approximate IC50 concentrations of 0.5 mg/L, 42 mg/L and 7,169 mg/L were calculated for MG, MGc and MGp, respectively (see IC50 determination below). The plasmid furthermore contains a *gfp* gene under the control of an inducible L-arabinose promoter and is maintained on average at 19 copies per bacterium. All strains were kindly provided by Prof. Alvaro San Millan (Centro Nacional de Biotecnología – CSIC, Madrid). The construction of MGc and MGp is described in detail in San Millan et al., 2016^18^. For the single-gene knockout mutant experiments, we used mutants Δ*oppA-D* from the Keio collection^47^.

### Growth conditions and media

*E. coli* overnight cultures were grown either in 8 mL lysogeny broth (LB; dose-response curves, competition experiments and experimental evolution assay) in 50 mL tubes or in 200 µL LB in 96-well plates (phenotypic growth assays) at 37 °C and 170 rpm agitation. Cultures grown overnight in tubes were washed twice with 0.8% NaCl and adjusted to an optical density at 600 nm (OD_600_) of 0.1. Cultures from well plates were used directly for experiments. The iron-limited casamino acid (CAA) medium (1% casamino acids, 5 mM K_2_HPO_4_ * 3H_2_O, 1 mM MgSO_4_*7 H_2_O, 25 mM HEPES buffer) was used for all experiments. Ceftazidime pentahydrate stocks of 1 mg/mL, ampicillin sodium salt stocks of 10 mg/mL, and pyoverdine stocks of 30 mg/mL were prepared in CAA. We used crudely purified pyoverdines from supernatants of the environmental *Pseudomonas* strain 3G07 (described in detail elsewhere^12^). Briefly, we grew the strain in 500 mL CAA medium, supplemented with 250 µM of the strong iron-chelator 2,2’-bipyridyl for 120 hours at 28

°C and shaken at 170 rpm. We then centrifuged the cultures (15,049 *g*) for 15 minutes, decanted the supernatant and adjusted its pH to 6 using 1 M HCl. We harvested the pyoverdines by running the supernatant over Amberlite XAD 16-N resins and eluting the pyoverdines with 50 % methanol. We lyophilized fractions containing the highest amount of pyoverdines (measured by fluorescence, excitation: 400 nm and emission: 460 nm) and stored it at −20 °C. Since we used crude pyoverdine extracts, the absolute concentration of pyoverdine was unknown and thus expressed relative to the highest concentration of 6 mg/mL used. All chemicals were purchased from Sigma-Aldrich, Buchs SG, Switzerland.

### Dose-response curves and IC50 calculation

To determine the inhibitory potential of pyoverdine, ceftazidime and ampicillin, we subjected the three *E. coli* strains to serially diluted crude pyoverdine extracts (highest concentration 6 mg/mL), to ceftazidime (highest concentration 4 mg/L) or to ampicillin (highest concentration 8 mg/mL (MG), 1024 mg/mL (MGc), 16,384 mg/mL (MGp)). To this end, bacterial cultures were grown overnight and prepared as described above. 2 µL of diluted culture were then added to a total of 200 µL medium with treatment on a 96-well plate in triplicates. Bacteria grown in CAA without treatment were included in each assay. Plates were incubated at 37 °C in a plate reader and the OD_600_ was measured every 15 minutes for 48 hours. We subsequently subtracted the blank and background values caused by the medium and the treatment from the growth values and calculated the area under the growth curve (AUC; integral) using the R package Growthcurver^53^. AUC values were then expressed relative to the untreated controls and plotted for each concentration. Finally, we fitted 5-parameter logistic regressions using the nplr package^54^ and extracted the IC50 values.

### Competitive fitness assays

To assess the potential cost of carrying antibiotics resistance on plasmids, we evaluated growth of MGp alone or in competition against MG and MGc in the absence of any treatment or in 0.5 mg/L ampicillin. Bacteria were cultured and prepared as described above. Competitions were initiated with a 1:1 mixture and were incubated alongside monocultures at 37 °C and shaken at 170 rpm for 46 hours. Prior to and after incubation, strain abundance was determined by flow cytometry. For this purpose, bacterial cultures were diluted in 1× phosphate buffer saline (PBS; Gibco, ThermoFisher, Zurich, Switzerland) and frequencies were measured using a Cytek Aurora 5L spectral analyzer (Cytek Biosciences, Amsterdam, The Netherlands) at a low flow rate and maximum mixing speed (1,500 rpm) for 2 seconds prior to measurement at the Cytometry Facility of the University of Zurich. To distinguish MGp from the competitors, we induced GFP expression by adding 0.2% L-arabinose 2 hours before each measurement (Laser: 488 nm, filter: 530/30;). Dead cells were stained with propidium iodide (PI) stain (2 μL of 0.5 mg/mL solution) (Laser: 355 nm, filter: 720/29). Before the competitions, we recorded 50,000 events, and after the competitions, we used a high-throughput plate loader system to record all events in a 5 µL volume.

For analysis, we used the software FlowJo (BD Biosciences, Ashland, OR) and followed the same gating strategy for all samples. First, we used forward- and side-scatter height values to separate bacterial cells from background. Within this gate, we excluded any doublings and retained only single cells using the forward- and side-scatter height and area values, respectively. Next, dead cells were excluded based on PI staining. Finally, we distinguished MGp from competitors by dividing cells into GFP-positive and negative populations. The relative fitness of MGp was then calculated using the formula ln(v) = ln{[a_48_×(1−a_0_)]/[a_0_×(1−a_48_)]}, where a_0_ and a_48_ are the frequencies of MGp at the beginning and at the end of the competition, respectively^55^. For each competition and treatment, a total of six replicates were measured, and two independent experiments were performed.

### Synergy degree of pyoverdine-antibiotic combination treatment

We used the Bliss independence model to calculate the degree of synergy (S) for growth of each *E. coli* strain in the pyoverdine-antibiotic combination treatment^23^. We used the formula *s* = *f*(*x*, 0) × *f*(0, *y*) − *f*(, *xy*), where*f*(*x*, 0) is the growth (integral) measured under antibiotic exposure at concentration X; *f*(0, *y*) is the growth (integral) measured under pyoverdine exposure at concentration Y; and*f*(, *xy*) is the growth (integral) measured in the combined treatment at concentration X and Y. If S = 0, pyoverdine and antibiotic act independently; S > 0 represents synergy, while S < 0 indicates antagonism.

### Experimental evolution experiment

To determine whether pyoverdine-antibiotic combination treatment can reduce selection for resistance, we experimentally evolved the three *E. coli* strains in single pyoverdine and ceftazidime treatments, in the combination of both and in a no-drug control treatment for 15 transfers. We used rounded values of the calculated IC50 concentrations for the single treatments (relative pyoverdine concentration of 0.16 and 0.4 mg/L for ceftazidime) and combined these concentrations for the combination treatment. We initiated the experiment with six independently evolving lineages per strain and treatment and arranged them diagonally in six 96-well plates. Evolving populations were surrounded by blank wells to reduce the risk of cross-contamination during transfer. Prior to the assay, strains were grown and prepared as described above, and 2 µL of culture were added to a total of 200 µL medium with or without antibacterial treatment. Plates were incubated at 37 °C at 170 rpm for 46.5 hours before bacterial cultures were diluted 100-fold into fresh treatment. At the end of each cycle, we added glycerol to the old plates at a final concentration of 15% and stored the plates at −80 °C.

### Phenotypic characterization of evolved populations

We investigated possible resistance evolution after repeated pyoverdine and ceftazidime treatment by comparing the growth of the evolved populations with that of the ancestor. To do so, we prepared overnight cultures of the evolved populations in 96-well plates as described above and then diluted the cultures 1000-fold into fresh treatment. We included six ancestors per strain and treatment, and 18 ancestors in plain medium. Plates were then incubated at 37 °C in a plate reader and growth was measured every 15 minutes for 48 hours. For analysis, we first subtracted the blank and background values from the growth data. Next, we calculated the AUC and expressed the values relative to the mean ancestor growth in plain medium. Finally, we calculated the growth difference between the evolved and the ancestors in the same treatment, which allows us to assess the growth gain of the evolved populations compared with the un-evolved populations. To determine whether pyoverdine reduced the selection for ceftazidime resistance in the combo treatment, MGp populations that had evolved in the combo treatment were subjected to conditions without pyoverdine, as well as to single treatments of pyoverdine and ceftazidime, and to the combo treatment, following the same procedure described above.

### Genomic analyses of evolved populations

For genomic analyses, we selected the four evolved populations with the highest growth gains observed in our phenotypic screen (Fig. 3) for each of the three treatments (pyoverdine, ceftazidime, combo) for strains MG and MGp. We further included 3 medium-adapted control populations and one ancestor clone per strain. For sequencing, we grew the populations and clones in 12 mL LB at 37 °C and 170 rpm and measured their OD_600_ after 6-8 hours. When cultures reached an OD_600_ between 0.8-1, we pelleted the cultures by centrifugation (7,500 *g*, 3 minutes), washed them in 0.8% NaCl, centrifuged again and finally resuspended the pellet in DNA shield buffer (Zymo research). Cultures were then sent to MicrobesNG (Birmingham, United Kingdom) for library preparation and whole-genome sequencing on the Illumina NovaSeq6000 platform (paired-end, 150 base-pair reads, minimum coverage 30X). Adapter sequences were trimmed using Trimmomatic v0.30^56^ with quality cut-off of Q15, reads were aligned to the closest available reference genome using BWA mem^57^, de novo assemblies were performed using SPAdes v3.7^58^ and contigs were annotated with Prokka v1.11^59^. Variants were predicted by using the breseq 0.37.0 pipeline^60,61^ using the polymorphism mode and the *E. coli* MG1655 reference genome (NC_000913.3) and pBGT plasmid genome^18^. Variants that were present in the ancestral clones relative to the reference sequence were excluded.

### Statistical analysis

All statistical analyses were performed in R 4.0.2^62^ and RStudio version 1.3.1056^63^. One-sample t-tests were used to compare the relative fitness and the degree of synergy in growth. We used analysis of variance (ANOVA) to test whether evolved populations grew significantly different than the ancestors and adjusted the p-values for multiple comparisons using the Tukey HSD test. The same analysis was applied to the experiments involving single-gene knockout mutants.

