## supporting_information for "Pyoverdine-antibiotic combination treatment: its efficacy and effects on resistance evolution in *Escherichia coli*"

This file contains supporting information for the article

**This supporting information contains:**

- 7 supporting figures S1 – S7
- 2 supporting tables S1 and S2

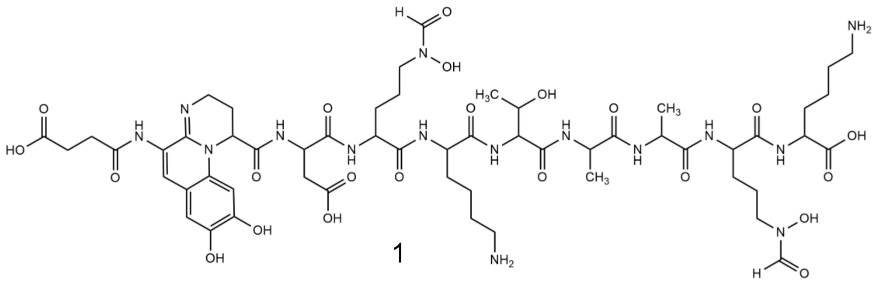

1

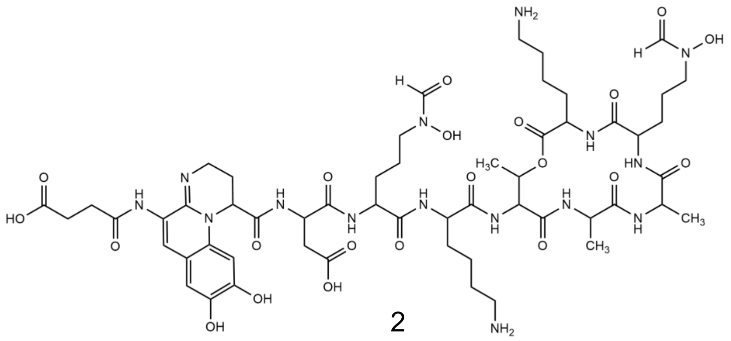

2

**Figure S1. Chemical structures of pyoverdine 3G07 from Vollenweider et al[1].** The chemical structures of pyoverdine 3G07 was elucidated using UHPLC-HR-MS/MS and revealed both a linear (1) and cyclic (2) form.

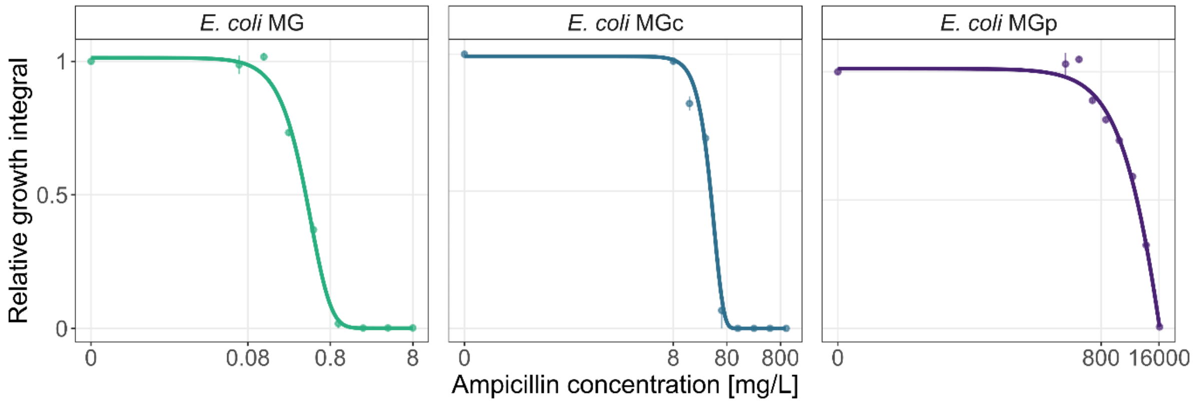

**Figure S2. Dose-response curves of *E. coli* strains MG, MGc and MGp after treatment with ampicillin.** We exposed the three strains to increasing concentrations of ampicillin and calculated the integral below the growth curves. The values were then scaled relative to the untreated control in CAA medium. Dots and error bars show mean values and standard error, respectively, across a minimum of six replicates per concentration. Dose-response curves were fitted using 5-parameter logistic regressions.

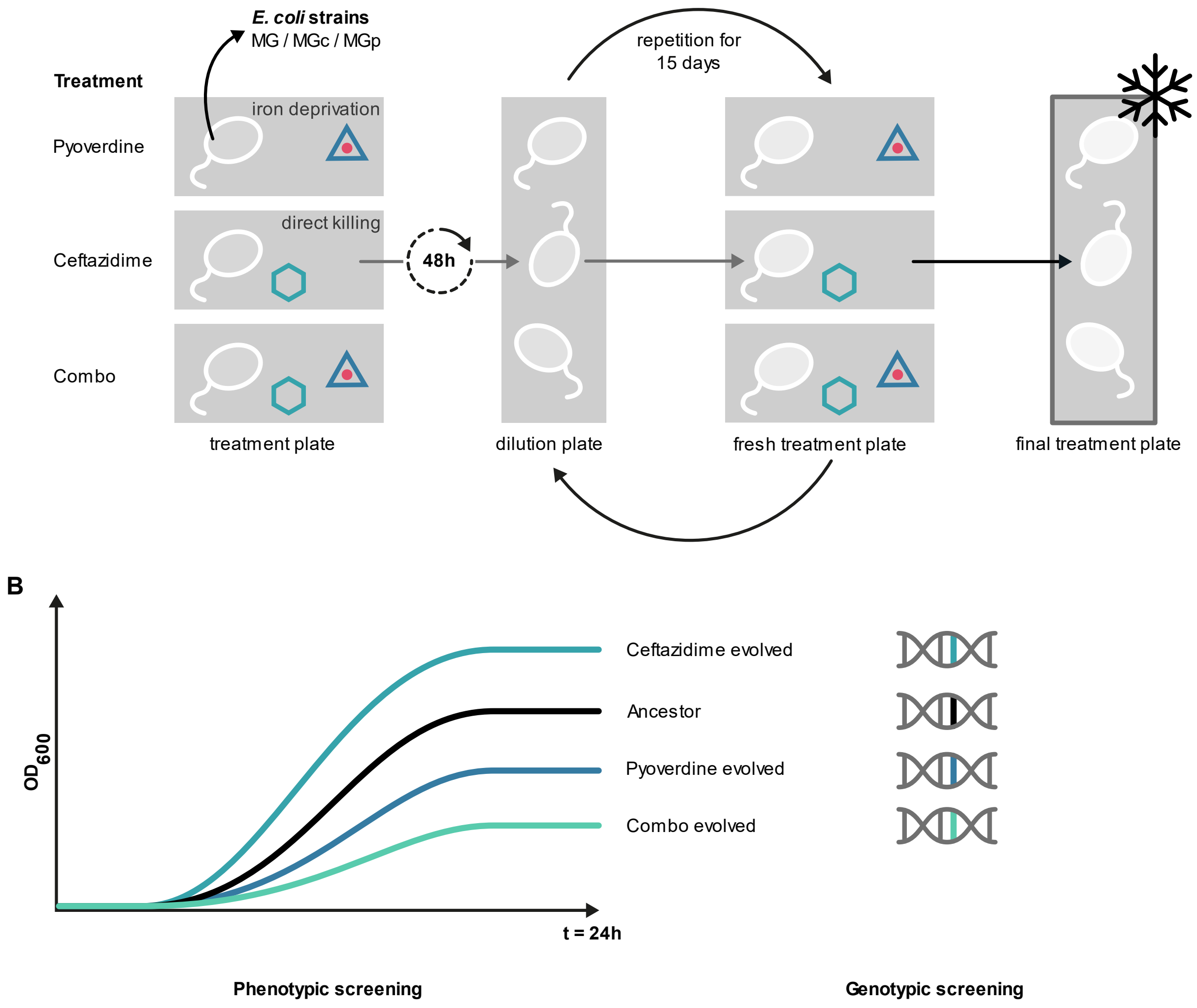

**Figure S3. Workflow of the experimental evolution experiment with MG, MGc and MGp and subsequent phenotypic and genotypic analyses.** (A) The assay involves treatments with pyoverdine, ceftazidime and their combination, as well as a no drug treatment control. We included six independently evolving lineages per treatment and strain and transferred cultures to fresh media every other day for a total of 15 transfers. The plate of the final time point was frozen as glycerol stocks at -80 °C. (B) Following experimental evolution, we compared the growth of the evolved lineages with the ancestors and sequenced 15 populations and one ancestor clone for both MG and MGp for in-depth genetic characterization.

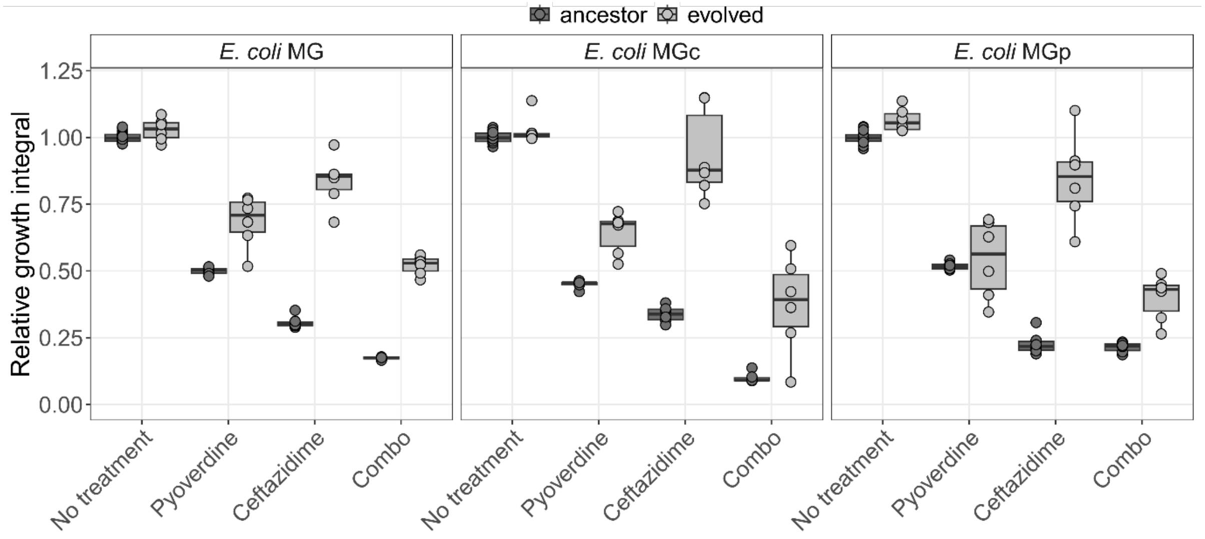

**Figure S4. Relative growth integral of evolved and ancestor *E. coli* MG, MGc and MGp populations after evolution in no-treatment, or the antibacterial treatments pyoverdine, ceftazidime or their combination.** We exposed evolved and ancestor *E. coli* populations to the treatment in which they evolved and quantified their growth by calculating the integral under the growth curves. Growth values were then scaled relative to the ancestor in plain CAA medium. The dots show mean values across 2 – 4 replicates from independent experiments. Significance levels are based on ANOVAS with adjusted p-values using the Tukey HSD method: * $p < 0.05$; ** $p < 0.01$; *** $p < 0.001$.

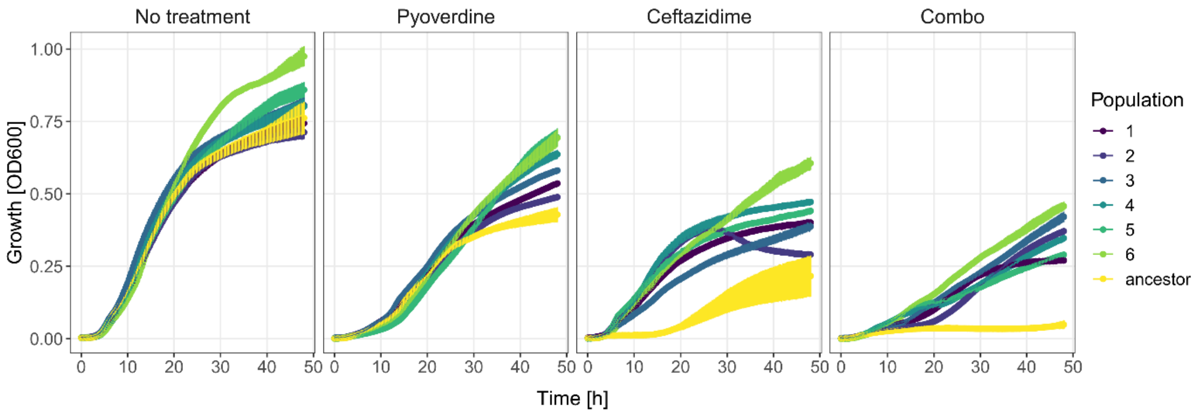

**Figure S5. Growth kinetics of *E. coli* MGp populations evolved in the combo treatment subjected to the no-treatment control, single pyoverdine 3G07 or ceftazidime treatments, or the combination treatment in which they evolved.** We exposed MGp populations that evolved in the combo treatment to single pyoverdine and ceftazidime treatments, as well as to plain medium (control) and the treatment they evolved in (combo) and monitored growth for 48 hours. Dots and error bars show mean values and standard error, respectively, across six replicates from two independent experiments.

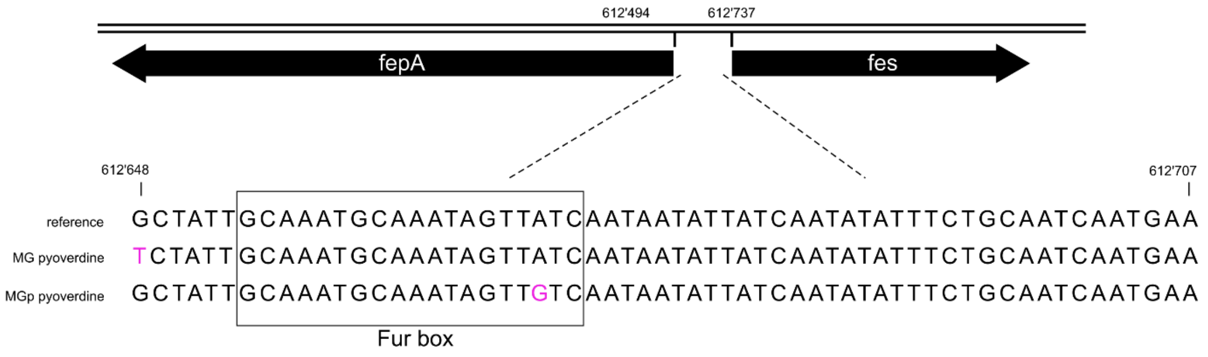

**Figure S6. Schematic representation of *fepA* and *fes*.** MG and MGp evolved in pyoverdine exhibited mutations in the intergenic region between *fepA*/*fes*.

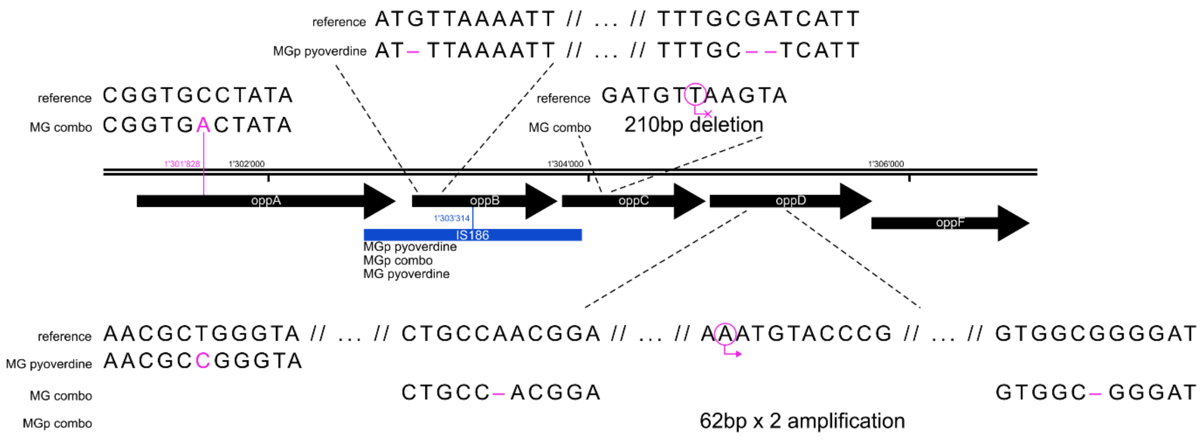

**Figure S7. Schematic overview of the *oppABCDF* operon**. Whole-genome sequencing revealed SNPs, deletions and amplifications in *oppA*, *oppB*, *oppC* and *oppD* genes after treatments containing pyoverdine.

### Table S1. Mutations frequencies of evolved *E. coli* MG and MGp populations.

| | | | LOW FREQUENCY | | HIGH FREQUENCY | |
|---|---|---|---|---|---|---|
| | | | MG | MGp | MG | MGp |
| SNPs | intergenic | control | 36 | 108 | 2 | 17 |
| | | pyoverdine | 27 | 113 | 1 | 22 |
| | | ceftazidime | 49 | 82 | | 20 |
| | | combo | 57 | 124 | 1 | 25 |
| | noncoding | control | 6 | 14 | | |
| | | pyoverdine | 5 | 14 | | |
| | | ceftazidime | 3 | 8 | | |
| | | combo | 10 | 8 | | |
| | nonsense | control | 1 | | | |
| | | pyoverdine | 1 | 4 | | 1 |
| | | ceftazidime | | | 2 | |
| | | combo | 1 | 2 | | 1 |
| | nonsynonymous | control | 17 | 17 | | 1 |
| | | pyoverdine | 14 | 11 | 5 | 1 |
| | | ceftazidime | 18 | 19 | 3 | 4 |
| | | combo | 18 | 20 | 7 | 5 |
| | pseudogene | control | 11 | 60 | | |
| | | pyoverdine | 21 | 62 | | 1 |
| | | ceftazidime | 18 | 70 | | |
| | | combo | 17 | 83 | | |
| | synonymous | control | 7 | 15 | | 3 |
| | | pyoverdine | 5 | 14 | | 4 |
| | | ceftazidime | 15 | 17 | | 5 |
| | | combo | 11 | 12 | | 4 |
| small deletions | coding | control | 1 | 1 | | |
| | | pyoverdine | 1 | 3 | 1 | 3 |
| | | ceftazidime | 2 | 1 | | 2 |
| | | combo | 3 | 1 | 3 | 2 |
| | intergenic | control | 2 | 8 | 1 | 6 |
| | | pyoverdine | | 2 | | 8 |
| | | ceftazidime | 1 | 8 | | 9 |
| | | combo | 1 | 9 | 1 | 8 |
| large deletions | | control | 1 | 4 | 1 | 9 |
| | | pyoverdine | | 4 | 1 | 16 |
| | | ceftazidime | | 2 | | 14 |
| | | combo | 2 | 2 | | 12 |
| small insertions | coding | control | 1 | | | 3 |
| | | pyoverdine | | 7 | | 4 |
| | | ceftazidime | 2 | 7 | 1 | 6 |
| | | combo | | | | 4 |
| | intergenic | control | 3 | 5 | | 6 |
| | | pyoverdine | 1 | 3 | | 8 |
| | | ceftazidime | 2 | 1 | | 9 |
| | | combo | | 4 | | 9 |
| | noncoding | control | | | | |
| | | pyoverdine | | | | |
| | | ceftazidime | | | | |
| | | combo | 1 | 1 | | |
| mobile element insertion | coding | control | 1 | | | |
| | | pyoverdine | 3 | 3 | 4 | 3 |
| | | ceftazidime | 2 | 2 | 2 | |
| | | combo | 5 | 3 | 1 | 4 |
| | intergenic | control | 1 | 1 | 1 | |
| | | pyoverdine | 6 | 1 | | |
| | | ceftazidime | | | 3 | |
| | | combo | | 1 | 2 | |
| large amplifications | | control | | | | |
| | | pyoverdine | | | | |
| | | ceftazidime | | | | |
| | | combo | | | | 1 |

### Table S2. Mutation type and position in the *opp* operon of *E. coli* MG and MGp.

| strain | treatment | population | gene name° | freq. | start position | gene strand | mut. category^Δ | ref seq | new seq | inactivated |
|---|---|---|---|---|---|---|---|---|---|---|
| MG | Pyoverdine | 1 | oppB | 0.85 | 1303314 | -1 | MOB | TTGTTA | | yes |
| MG | Pyoverdine | 2 | oppB | 0.73 | 1303314 | 1 | MOB | TTGTTA | | yes |
| MG | Pyoverdine | 5 | oppB | 0.67 | 1303314 | -1 | MOB | TTGTTA | | yes |
| MG | Pyoverdine | 5 | oppB | 0.74 | 1303314 | 1 | MOB | TTGTTA | | yes |
| MG | Pyoverdine | 6 | oppD | 1 | 1304908 | NA | nonsyn. SNP | T | C | |
| MG | Combo | 1 | oppD | 1 | 1305452 | NA | small deletion | G | | yes |
| MG | Combo | 3 | oppD | 1 | 1304979 | NA | small deletion | A | | yes |
| MG | Combo | 5 | oppA | 0.65 | 1301828 | NA | nonsyn. SNP | C | A | |
| MGp | Pyoverdine | 4 | oppC | 1 | 1303841 | NA | large deletion | 210-bp | | yes |
| MGp | Pyoverdine | 3 | oppB | 0.37 | 1302901 | NA | small deletion | G | | yes |
| MGp | Pyoverdine | 2 | oppB | 0.51 | 1303363 | NA | small deletion | G | | yes |
| MGp | Pyoverdine | 2 | oppB | 0.51 | 1303364 | NA | small deletion | A | | yes |
| MGp | Pyoverdine | 1 | oppB | 1 | 1303314 | -1 | MOB | TTGTTA | | yes |
| MGp | Combo | 5 | oppD | 0.84 | 1305242 | NA | large amplification | 62-bp | | |
| MGp | Combo | 3 | oppB | 1 | 1303314 | -1 | MOB | TTGTTA | | yes |
| MGp | Combo | 3 | oppB | 1 | 1303314 | 1 | MOB | TTGTTA | | yes |
| MGp | Combo | 4 | oppB | 1 | 1303314 | -1 | MOB | TTGTTA | | yes |

° oppA oligopeptide ABC transporter periplasmic binding protein
oppB murein tripeptide ABC transporter/oligopeptide ABC transporter inner membrane subunit OppB
oppC murein tripeptide ABC transporter/oligopeptide ABC transporter inner membrane subunit OppC
oppD murein tripeptide ABC transporter/oligopeptide ABC transporter ATP binding subunit OppD

Δ MOB: mobile element insertion (IS186); nonsyn. SNP: nonsynonymous single nucleotide polymorphism
